# A polarized genome is required for fitness of the obligate intracellular, tick-borne pathogen *Rickettsia parkeri*

**DOI:** 10.64898/2026.11.11.765969

**Authors:** Trung Nguyen, Andrew C. Krusenstjerna, Mathilde Gonin, Wanda M. Figueroa-Cuilan, Erika L. Smith, Aayush Sapkota, Himaja Maddali, Erin D. Goley

## Abstract

Alphaproteobacteria of the order Rickettsiales include biomedically important, yet understudied arthropod-borne human pathogens. These pathogens lack obvious morphological asymmetry, but encode homologs of key mediators of polar processes in the model Alphaproteobacterium *Caulobacter crescentus*. These include the polar organizing protein PopZ, the chromosome segregation proteins ParA and ParB, and the spatial regulator of cell division MipZ. Motivated by this observation, we used the tick-borne human pathogen *Rickettsia parkeri* to ask if the chromosome is polarized and coordinated with cell division in this group. Consistent with that hypothesis, we found that transposon insertion mutants in *parA*, *parB*, *mipZ*, or *popZ* resulted in cell division defects. Fluorescence imaging revealed cell cycle-dependent unipolar-to-bipolar localization dynamics of ParB and MipZ, and pole-to-midcell dynamics of the division protein ZapA. This patterning parallels the model of chromosome segregation and cell division site selection established in *Caulobacter*. Unlike its unipolar-to-bipolar dynamics in *Caulobacter*, however, PopZ remained at a single cell pole throughout the cell cycle in *R. parkeri.* PopZ maintained its direct interaction with ParA and ParB. Interestingly, disruption of *parA* or *parB*, but not *popZ or mipZ*, caused a small plaque phenotype, suggesting a defect in growth or spread in a monolayer of host cells in the absence of proper chromosome segregation. We propose that genome polarization in *R. parkeri* and other rickettsiae is critical both for fundamental cellular processes and for interactions with host cells that promote fitness in their infectious niche.

**IMPORTANCE:** Bacterial species in the order Rickettsiales encompass globally distributed arthropod-borne pathogens that elicit potentially deadly infections. Despite their importance, the obligate intracellular lifestyle of rickettsial species imposes significant technical challenges to their study. Here, we apply recently developed imaging and genetic tools to probe the regulation of *Rickettsia parkeri*’s genome dynamics and cell division. Our findings address an important knowledge gap regarding the molecular mechanisms underlying fundamental cell biological processes in rickettsial pathogens. These mechanisms impact bacterial physiology, pathogenesis, and virulence, warranting a thorough understanding to inform therapeutic interventions that prevent rickettsial proliferation and infection progression.

## INTRODUCTION

Alphaproteoacteria of the order Rickettsiales are notable for their strict association with eukaryotic host cells. Evolutionary adaptation to their host environment has led to genome reduction in the Rickettsiales, leaving only a portion of the proteome compared with that of related free-living species [1]. Within this order, the genus *Rickettsia* includes arthropod-borne human pathogens that cause infections worldwide and collectively impose a considerable public health burden [2,3]. These include members of the Spotted Fever Group (SFG), etiologic agents of human tick-borne diseases ranging from mild and self-limiting (*Rickettsia parkeri* rickettsioses caused by *R. parkeri*) to potentially lethal (Rocky Mountain Spotted Fever caused by *R. rickettsii*) [2,3]. Over the past two decades, the United States has witnessed a surge in the reported cases of SFG rickettsial infections, with the Centers for Disease Control and Prevention declaring SFG bacteria to be emerging pathogens [2–4]. Despite their biomedical importance, their obligate intracellular lifestyle complicates investigation of their biology, with a lack of axenic culture and limitations on genetic manipulation hindering progress.

*R. parkeri* causes a relatively mild infection in humans, but possesses pathogenic, physiological, and genetic similarities to more virulent SFG and rickettsial species. It therefore represents a tractable model for probing rickettsial cell biology [5]. Like other SFG pathogens, the infectious cycle of *R. parkeri* starts with attachment and internalization of the pathogen into the host cell, rapid escape from the phagosome, and growth in the host cytoplasm. Actin-based motility of *R. parkeri* mediates dissemination into neighboring host cells and to distal tissues in an animal host [6]. Although the obligate intracellular nature of *Rickettsia* species complicates their study, progress has been made on this front for *R. parkeri* in recent years. A collection of ∼200 transposon insertions in ∼110 different genes (out of ∼1350 total protein-coding genes) has laid the foundation for mechanistic studies of *R. parkeri* pathogenesis and host interactions [7,8]. In addition, our laboratory has established high-throughput, quantitative live-imaging techniques for assessment of *R. parkeri* growth dynamics and morphogenesis inside host cells [9].

In this study, we leverage these technical advances to investigate the molecular mechanisms underlying chromosome duplication and segregation and their coordination with cytokinesis – processes that must be tightly coordinated for successful replication. These fundamental aspects of bacterial cell biology have been extensively studied in free-living organisms, including the alphaproteobacterial model *Caulobacter crescentus* [10–13]. The site of cell division is marked by the tubulin homolog FtsZ, which polymerizes into a dynamic ring (Z-ring) and serves as a platform for assembly of the divisome that drives cell constriction. ZapA is a conserved coiled-coil protein that interacts directly with FtsZ and plays an important role in condensation of the Z-ring for efficient constriction [14,15]. In this study, we use *R. parkeri* ZapA as a proxy to visualize the site of Z-ring assembly and cell division.

In *C. crescentus*, chromosome organization and division site selection are tightly coordinated in time and space. *C. crescentus* deploys the ParABS system to segregate its chromosome [16,17]. In this system, the CTPase ParB binds a chromosomal locus called *parS*, which is located near the chromosomal origin of replication (*oriC*), forming a ParB-*parS* partition complex [17]. In *C. crescentus*, ParB-*parS* and *oriC* are anchored at the old cell pole via direct interaction of ParB with the polar organizing protein PopZ [18,19]. A second ParB-*parS* complex is formed at this pole shortly after *parS* is replicated. A third component of the system, the ATPase ParA, forms a dimer upon binding ATP and binds the nucleoid in a sequence-nonspecific manner [20,21]. After replication, one ParB-*parS* complex interacts with DNA-bound ParA and stimulates its ATPase activity, releasing ParA from the chromosome. ParB-*parS* iteratively contacts new ATP-bound ParA dimers on the nucleoid, promoting their release and ultimately segregating ParB-*parS* to the opposite pole where the complex is captured by a newly accumulated PopZ condensate [22–24].

At the heart of the spatiotemporal checkpoint that coordinates cell division with DNA segregation lies a deviant Walker A-type ATPase called MipZ, which is an inhibitor of FtsZ polymerization [25,26]. MipZ directly associates with ParB and with DNA and, following ParB-*parS* translocation, establishes a bipolar gradient whose concentration peaks at the two cell poles and decreases toward midcell [25,26]. Since MipZ inhibits FtsZ polymerization, it effectively restricts divisome formation to midcell and coordinates division-site selection with chromosome segregation in time and space. Depletion of ParA, ParB, or MipZ, or deletion of *popZ,* all lead to filamentation in *C. crescentus*, reflecting their roles in establishing the site of division [25,26].

Polar targeting of the chromosome in *C. crescentus* is integrated with other polarized cellular events, culminating in the formation of distinct appendages at each pole, asymmetric cell division, and different cell-cycle statuses of the resulting daughters [27]. *R. parkeri* and other *Rickettsia* species are rod-shaped and lack apparent morphological asymmetry. Despite this lack of obvious polarity, they encode putative homologs of PopZ, ParA, ParB, and MipZ (**Figure 1**), suggesting they may establish intracellular polarity to coordinate key cell biological processes, as in *C. crescentus*.

**Figure 1.**
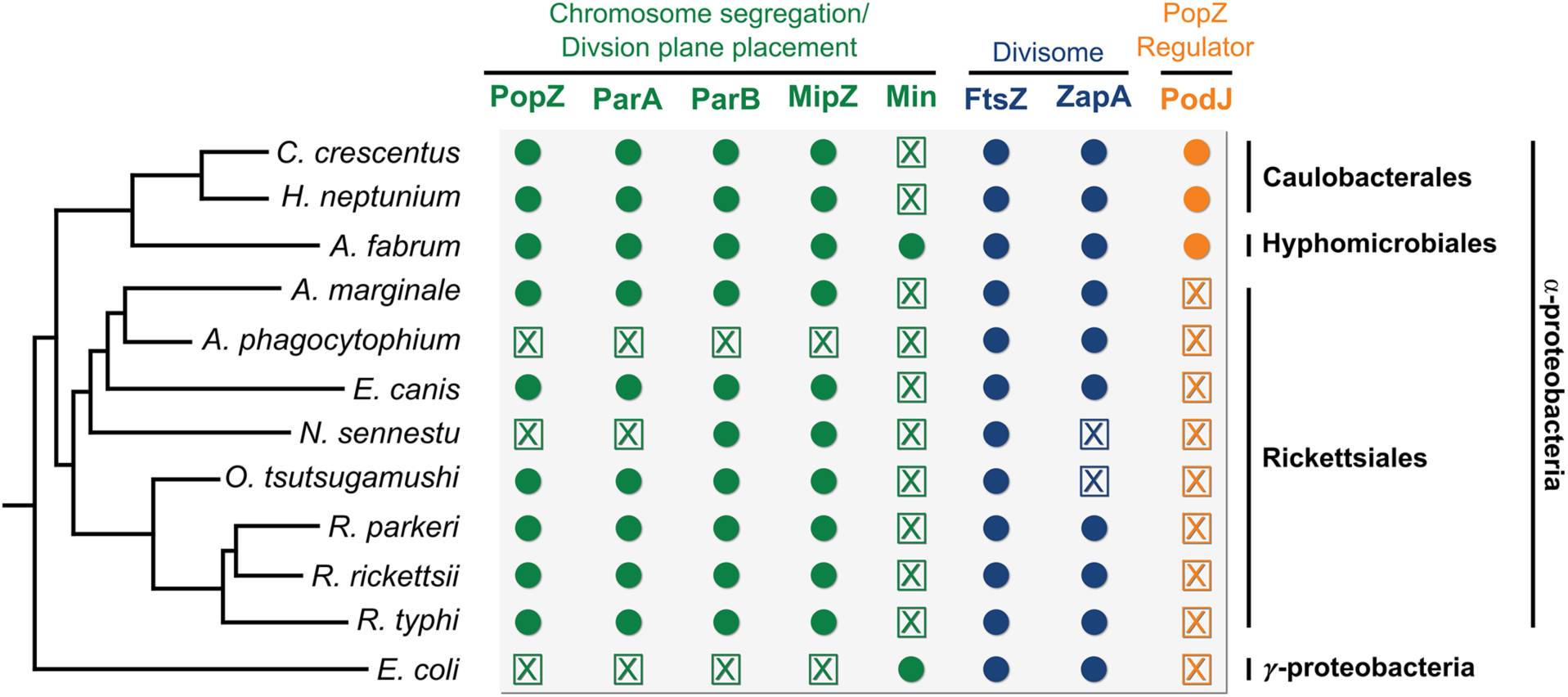
Conservation of chromosome segregation and cell division proteins in the Alphaproteobacteria. Phylogeny of diverse Alphaproteobacteria and the gammaproteobacterial representative *E. coli* was constructed using phyloT. Genes encoding chromosome segregation and cell division regulators (green), components of the divisome (blue), and the predicted PopZ regulator PodJ (orange) were identified using BioCyc [76]. Filled circles and crossed boxes denote the presence and absence of genes, respectively.

Here, we report that in *R. parkeri*, the mechanism of spatiotemporal coordination of chromosome dynamics with cell division is generally conserved relative to *C. crescentus*, except for distinct localization dynamics of PopZ. Perturbations in chromosome segregation, chromosome anchoring, or division site selection differentially impact *R. parkeri*’s ability to grow within host cells and spread within a monolayer. Notably, loss of chromosome segregation, but not chromosome anchoring or division site selection, significantly impairs growth and spread in a host monolayer. Our data demonstrate conservation of fundamental cell cycle mechanisms in Alphaproteobacteria with remarkably different lifestyles. Our findings also indicate that subcellular organization of *R. parkeri* is functionally linked to host interactions and fitness in the intracellular growth environment. These mechanisms may represent previously uncharacterized targets for infection management.

## RESULTS

### Putative chromosome segregation and cell division proteins are encoded in the *Rickettsiales*

As a first step in examining the cell biology of *Rickettsia*, we searched for putative regulators of chromosome segregation and cell division in the genomes of representative *Rickettsiales.* Our analysis included well-characterized species in the alphaproteobacterial orders *Caulobacterales* and *Hyphomicrobiales* for comparison purposes, as well as *Escherichia coli* as a representative outgroup. Except for *Anaplasma phagocytophilum* and *Neorickettsia sennetsu*, most members of the *Rickettsiales* encode putative homologs of the pole organizing protein PopZ, the chromosome segregation machinery ParAB, and the division regulator MipZ (**Figure 1**).

### Transposon insertions in *parAB*, *popZ*, or *mipZ* impair cell division

A transposon mutant in *parB* was previously isolated and reported to exhibit a small plaque phenotype in *R. parkeri*, suggesting this factor may be important for growth in host cells [7]. We fortuitously isolated additional transposon mutants in *popZ*, *mipZ*, and *parA* (**Figure 2A**), allowing us to pursue functional characterization of these factors in *R. parkeri*. Each transposon insertion cassette carries a fluorescent protein gene for quantitative imaging and an antibiotic resistance cassette for selection of transformants (**Figure 2A**).

**Figure 2.**
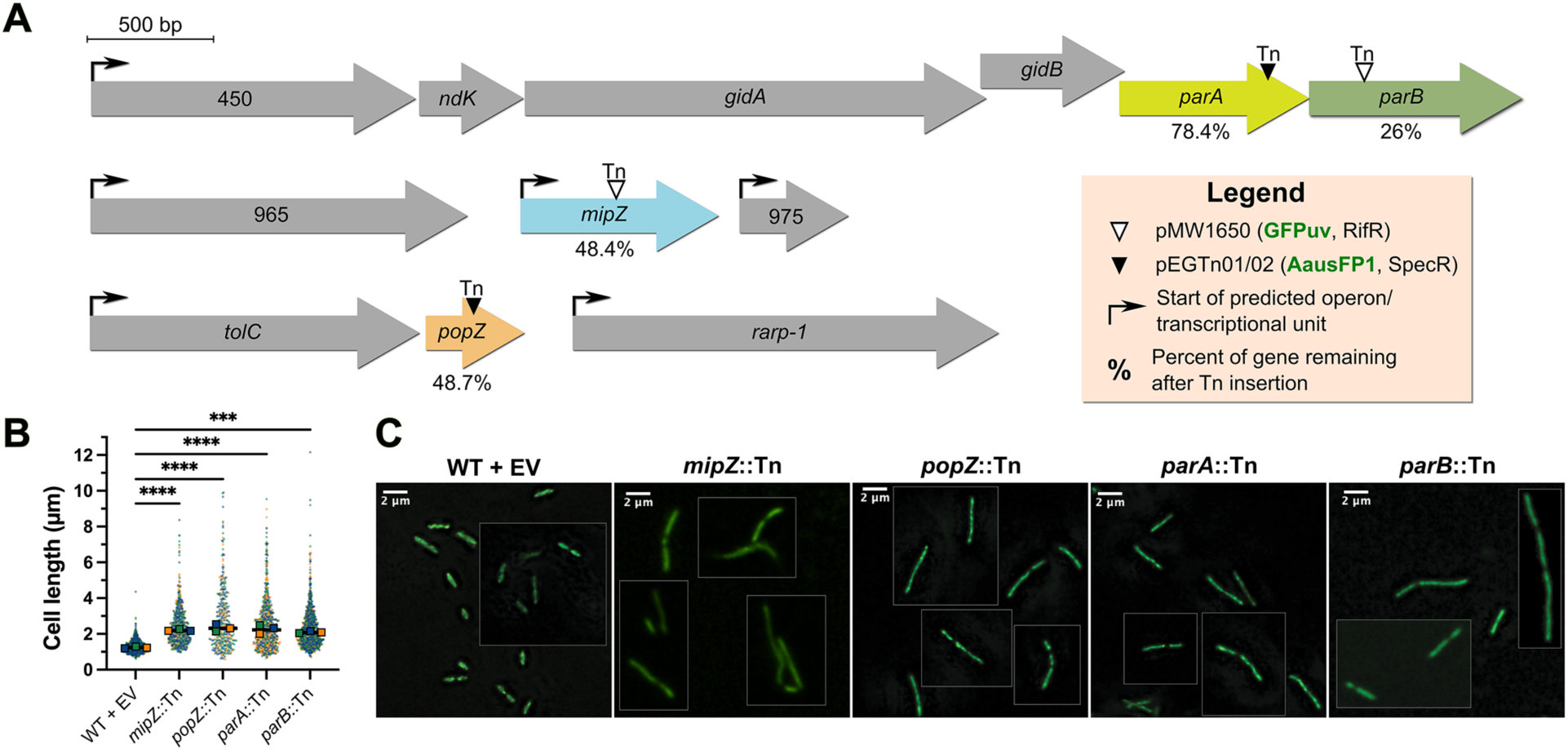
Transposon mutagenesis of chromosome segregation and cell division regulators results in filamentation and cell division defects. **A.** Map of the genomic loci for *parA*, *parB, mipZ,* and *popZ*, the transposon insertion site, and the percentage of the truncated proteins produced relative to their full-length version. Operon prediction was obstained using BioCyc [76]. **B.** Dot plots of cell length in MicrobeJ using cytoplasmic AausFP1 or GFPuv expressed from the chromosomal insertion of the transposon mutants compared to WT *R. parkeri* carrying an empty vector (EV) control. Circles represent individual cell measurements from three technical replicates (blue, orange, and green; >110 cells/replicate) and outlined boxes represent median values for each replicate. Lines indicate the mean of replicate medians. Ordinary one-way ANOVA analysis with Dunnett’s multiple comparison tests was performed to compare the population median values (n = 3) of each mutant to WT *R. parkeri* and determine *P* values (***, p = 0.0001, ****, p < 0.0001), assuming normal (Gaussian) distribution of the median length values. **C.** Composite images of live, intracellular *R. parkeri* transposon mutant cells in A549 host cells at ∼24 hpi. Boxed images represent different fields of view combined to increase the number of representative cells displayed. WT + EV = EGRp0035; *mipZ*::Tn = EGRp0022; *popZ*::Tn = EGRp0128; *parA*::Tn = EGRp0052; *parB*::Tn = EGRp0034.

We examined the site of insertion in each mutant to inform the putative effect of each on the encoded protein and found that each is likely to result in complete loss of function. In the *parA*::Tn mutant strain, the transposon insertion resulted in a truncation of ParA at residue 200 of 256 and ablated the C-terminal domain that is required for non-specific DNA binding [28]. *parA* and *parB* are the last two genes in a putative six-gene operon, so disruption of *parA* may also impact expression of *parB* (**Figure 2A**). In the *parB*::Tn mutant strain, the transposon insertion disrupts the entire putative DNA-binding region and C-terminal dimerization domain, leaving only a segment of the N-terminus predicted to be involved in nucleotide binding and protein-protein interaction (**Figure 2A**) [29]. The insertion in *mipZ*::Tn truncates MipZ at residue 128 of 266, ablating a region that mediates DNA binding, which is required for MipZ gradient formation (**Figure 2A**) [25,30]. Finally, in the *popZ*::Tn mutant strain, PopZ was truncated at residue 65 in the first C-terminal oligomerization helix and consequently is predicted to fail to form a condensate (**Figure 2A**) [18,19,31–33]. Different operon prediction algorithms yield distinct results on whether *popZ* is the last gene in a two-gene operon or if it is second in a four-gene operon also including the periplasmic protein RARP-1. However, our phenotypic characterization supports the former and suggests *popZ*::Tn does not result in polar effects on RARP-1 function (see below). We conclude that each of the above transposon mutants is likely to result in complete loss of function of the encoded protein, and that the *parA*::Tn mutant may also disrupt ParB production.

To determine potential effects of these mutations on cell division or other aspects of morphology, we imaged each strain at ∼24 hours post-infection (hpi) of A549 human lung epithelial cells using GFPuv or AausFP1 signal (expressed from the transposon) to delineate bacterial cell outlines. We found that each mutant exhibited a significant increase in cell length and misplaced constriction sites when compared to wild-type (WT) *R. parkeri* harboring an empty vector encoding cytoplasmic AausFP1 (**Figure 2B-C**). These phenotypes are consistent with a role for each of these proteins in chromosome segregation and/or cell division, as predicted from their functions in *C. crescentus*. These data also support our hypothesis that these are loss-of-function transposon mutants.

### Cell cycle dynamics of genome organization and cell division in *R. parkeri*

We next set out to characterize the spatiotemporal dynamics of chromosome segregation and division plane positioning in *R. parkeri* by visualizing fluorescent fusions to ParB, PopZ, and MipZ. ParB was tagged with mNeonGreen (ParB-mNG) and expressed from a replicating plasmid under the control of the predicted *tolC* promoter (comprising the 200 bp upstream of the *tolC* gene, which precedes *popZ* in a predicted operon – **Figure 2A**). This plasmid also produces cytoplasmic BFP for visualization of bacterial cells. Although we have access to the *parB*::Tn mutant strain, we have been unable to isolate stably transformed clones after electroporation of this strain with pRAM-based plasmids for unknown reasons. We therefore visualized ParB-mNG in a WT background, i.e., in the presence of native ParB. At ∼24 hpi of A549 cells, we observed *R. parkeri* cells with a single polar focus of ParB-mNG, cells with bipolar ParB-mNG, and cells with one polar and one non-polar ParB-mNG focus.

We conducted demograph analysis to understand the cell cycle dynamics of ParB-mNG. Demographs revealed that ParB-mNG was unipolar in newborn G1 cells, with a second focus appearing and moving across the cell body as the cell cycle progressed, reflecting *parS* segregation. Eventually, ParB-mNG exhibited a bipolar pattern in pre-divisional cells, indicating *parS* was partitioned to the two poles prior to division (**Figure 3A-B**). The expression of ParB-mNG did not alter cell length of this *R. parkeri* strain compared to WT *R. parkeri* producing cytoplasmic BFP from an empty vector (**Supplemental Figure S1A-B**), suggesting that mild overproduction of ParB-mNG did not disrupt its normal function.

**Figure 3.**
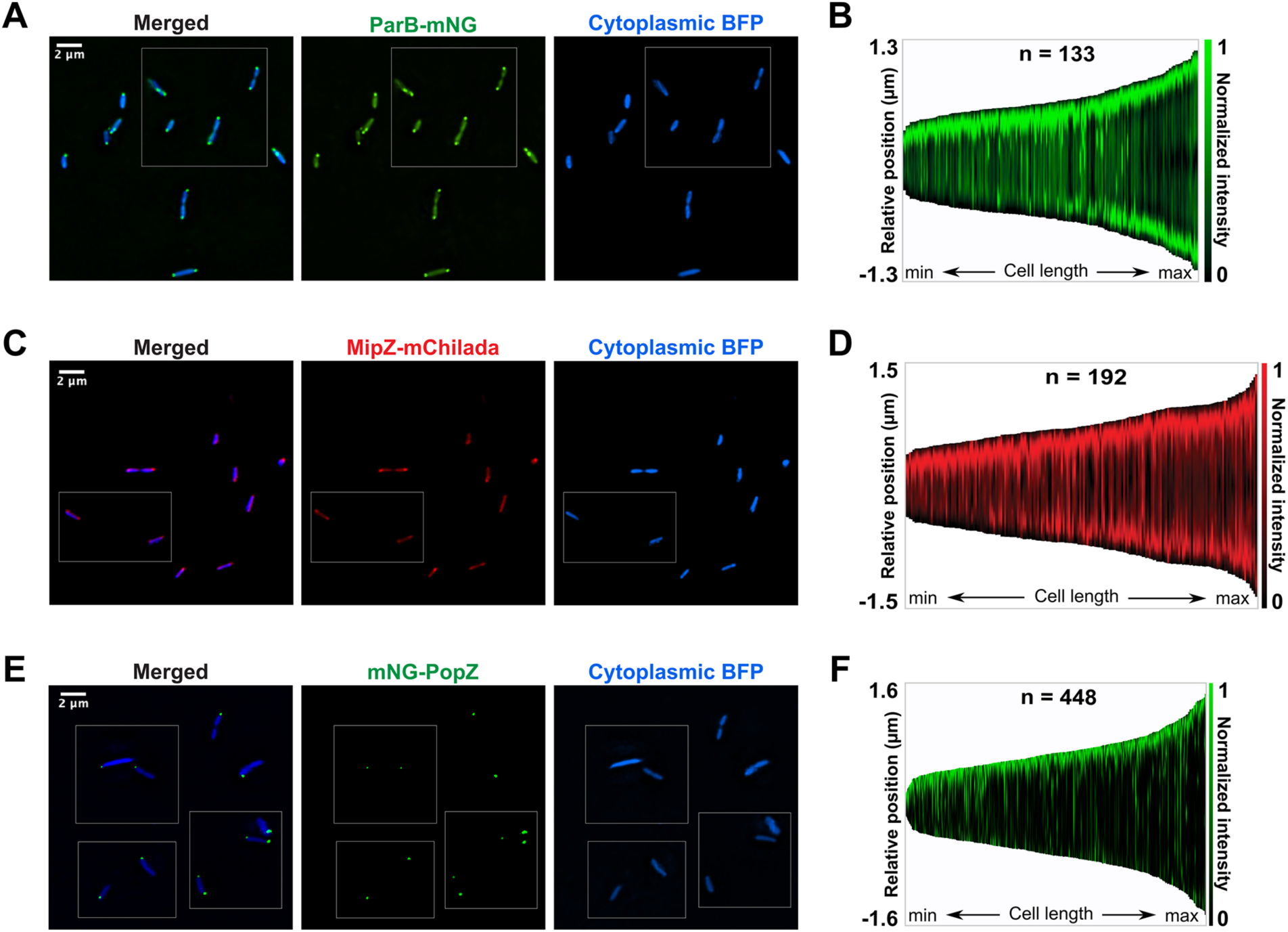
Subcellular localization dynamics of chromosome segregation and cell division factors in *R. parkeri*. Representative epifluorescence micrographs and demographs of live *R. parkeri* in A549 host cells at 24 hpi. *R. parkeri* expressed cytoplasmic *tagBFP* from the *ompA* promoter with (**A-B**) *parB-mNG* from the *tolC* promoter in a WT background (EGRp0015), (**C-D)** *mipZ-mChilada* from the *zapA* promoter in a *mipZ*::Tn background (EGRp0018), and (**E-F**) *mNG-popZ* from the *tolC* promoter in a WT background (EGRp0053). Boxed images represent different fields of view to increase the number of representative cells displayed. Demographs were generated by identifying cell outlines using TagBFP signal and plot mNG or mChilada signal as a function of cell length in MicrobeJ. Each demograph represents cells from one of three technical triplicates for each strain.

Next, we tagged MipZ with the red fluorescent protein mChilada under the control of its own promoter in a plasmid also producing cytoplasmic BFP to probe the inhibitory and permissive zones for Z-ring assembly. Consistent with the localization pattern of its predicted direct interactor ParB, demograph analysis showed that MipZ-mChilada expressed in a *mipZ*::Tn mutant background formed a gradient, with the most intense signal at a single cell pole in G1 cells and with intensity at both poles after entry to S phase. The lowest MipZ-mChilada concentration was at midcell as cells prepared to divide (**Figure 3C-D**). The MipZ-mChilada fusion restored WT cell length to the *mipZ*::Tn mutant, suggesting it is a functional fusion (**Supplemental Figure S1E-F**).

As we previously reported [9,34], ZapA-mChilada expressed from its own promoter in a pRAM-based plasmid in WT *R. parkeri* localizes at a single cell pole that is presumably opposite the pole bearing a MipZ focus in G1 cells and moves to midcell in pre-divisional cells (**Supplemental Figure S1C-D**). The localization dynamics of ParB, MipZ, and ZapA are consistent with conservation of the model established in *C. crescentus*.

Finally, we analyzed the subcellular localization of PopZ by tracking PopZ fused to mNeonGreen (mNG-PopZ) and expressed under the *tolC* promoter in a WT background. Surprisingly, and unlike *C. crescentus*, most bacteria (∼80%) exhibited a single polar focus of mNG-PopZ. Demograph analysis of mNG-PopZ revealed a unipolar localization of the protein throughout the cell cycle rather than unipolar-to-bipolar dynamics as observed for ParB-mNG and MipZ-mChilada (**Figure 3E-F**). Occasional bipolar localization of PopZ was observed (7% of cells), but there was no obvious correlation to cell cycle progression (**Figure 3F, Supplemental Figure S2**). PopZ from *C. crescentus* undergoes phase separation that is dependent on protein concentration [33]. mNG-PopZ was produced from a replicating plasmid in addition to native expression from the chromosomal locus in *R. parkeri.* Therefore, we hypothesize that the occasional appearance of more than one PopZ focus in a subset of cells (**Supplemental Figure S2**) is an artifact of PopZ exceeding its cellular saturation concentration, rather than a physiologically meaningful localization pattern. The mNG-PopZ fusion was functional, as it could restore WT length to the *popZ*::Tn mutant and did not disrupt cell length when produced in a WT background (**Figure 3E, Supplemental Figure S1G-H**). Collectively, our data show that while chromosome segregation dynamics and division plane positioning in *R. parkeri* are consistent with the mechanism established in *C. crescentus*, the unipolar localization of PopZ hints at distinctions in the mechanism of polar ParB-*parS* docking at the new cell pole by PopZ in *R. parkeri*.

### Direct interaction of *R. parkeri* PopZ with ParA and ParB

In *C. crescentus*, PopZ interacts directly with ParA and ParB to regulate chromosome segregation and polar *parS* targeting [19,32,35]. To test potential physical association between PopZ and ParA or ParB in *R. parkeri*, we leveraged *Escherichia coli* as a heterologous expression system. *E. coli* does not encode homologs of PopZ, ParA, or ParB (**Figure 1**), and prior work demonstrated that PopZ from *C. crescentus* forms a single polar condensate that can recruit its direct interaction partners when produced in *E. coli* [18,19,32,35]. We ectopically produced mCherry-PopZ from *R. parkeri* in *E. coli* and found that it formed a focus at the cell pole (**Supplemental Figure S3C-D**). Therefore, *E. coli* could serve as a heterologous system to examine direct interaction of *R. parkeri* PopZ with ParA and ParB.

When PopZ expression was not induced, ParA-msfGFP showed a nucleoid-associated pattern, as expected for a sequence-nonspecific DNA binding protein (**Figure 4A-B**). ParB-msfGFP exhibited a cytoplasmic localization, consistent with a lack of a *parS* binding sequence in *E. coli* (**Figure 4G-H**). Upon induction of mCherry-PopZ, ParB-msfGFP and ParA-msfGFP were each recruited to the mCherry-PopZ focus, indicating their physical association with the PopZ condensate (**Figure 4C-D** and **I-J**). As a control, msfGFP alone was evenly distributed in *E. coli* cytoplasm in the presence of mCherry-PopZ (**Supplemental Figure S3A-B**). We conclude that PopZ likely anchors ParB-*parS* at the cell pole *via* direct interaction with ParB, and that the PopZ-ParA interaction may contribute to function of ParA in chromosome segregation as in *C. crescentus* [16].

**Figure 4.**
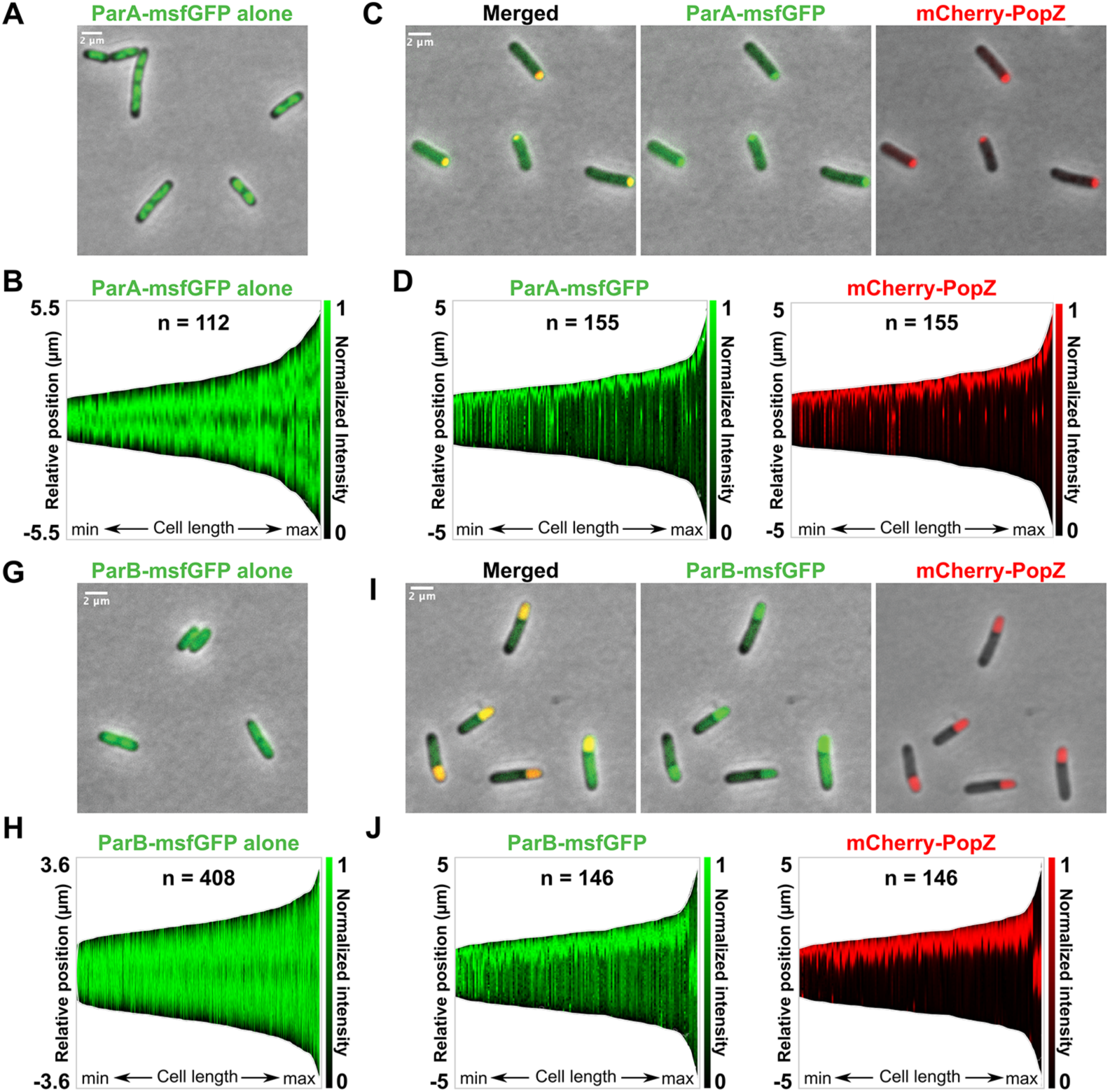
Direct interaction of *R. parkeri* PopZ with ParA and ParB in *E. coli*. Representative epifluorescence micrographs and demographs of *E. coli* BL21 cells producing *R. parkeri* fluorescently tagged proteins at 2 hours post-induction. **A-B.** *R. parkeri* ParA-msfGFP localized to the nucleoid when produced alone (pEG2201). **C-D.** Nearly all *E. coli* cells co-producing *R. parkeri* ParA-msfGFP with mCherry-PopZ (pEG2205) showed colocalization of ParA-msfGFP with the mCherry-PopZ foci. **G-H.** *R. parkeri* msfGFP-ParB signal was diffuse when produced alone (pEG2209). **I-J.** Co-expression with mCherry-PopZ resulted in recruitment of ParB-msfGFP to the mCherry-PopZ condensate in almost all *E. coli* cells (pEG2210). Representative demographs of one of three technical triplicates display msfGFP and mCherry signal as a function of cell length.

### Dysregulation of cell division in *popZ*::Tn and *mipZ*::Tn

So far, our data are consistent with the model that PopZ is required for polar localization of ParB, which then recruits MipZ to the pole. MipZ, in turn, positions the Z-ring. To directly test this model, we inspected MipZ-mChilada and ZapA-mChilada localizations in the *popZ*::Tn and *mipZ*::Tn mutant strains, respectively. In the *popZ*::Tn mutant, MipZ-mChilada formed one or more foci along the length of each filamentous cell body. A gradient of intensity was still observed around each peak of MipZ-mChilada fluorescence, suggesting MipZ’s biochemical activity and interaction with DNA was preserved while its localization was disrupted (**Figure 5A-B**). In *C. crescentus ΔpopZ* cells, DNA replication and segregation by ParABS continues, with multiple ParB-*parS* complexes segregated relatively evenly along the length of each filamentous cell but not anchored at the cell poles [18,19]. We observed a similar pattern for MipZ-mChilada in our *popZ*::Tn mutant, with relatively regularly spaced peaks of MipZ-mChilada intensity along the cell body. In the *mipZ*::Tn mutant, we observed multiple ZapA-mChilada foci along the filamentous cell body, consistent with misregulation of Z-ring placement (**Figure 5C-D**). These data provide further evidence that obligate intracellular rickettsial species follow a model similar to *C. crescentus* chromosome segregation and division site selection.

**Figure 5.**
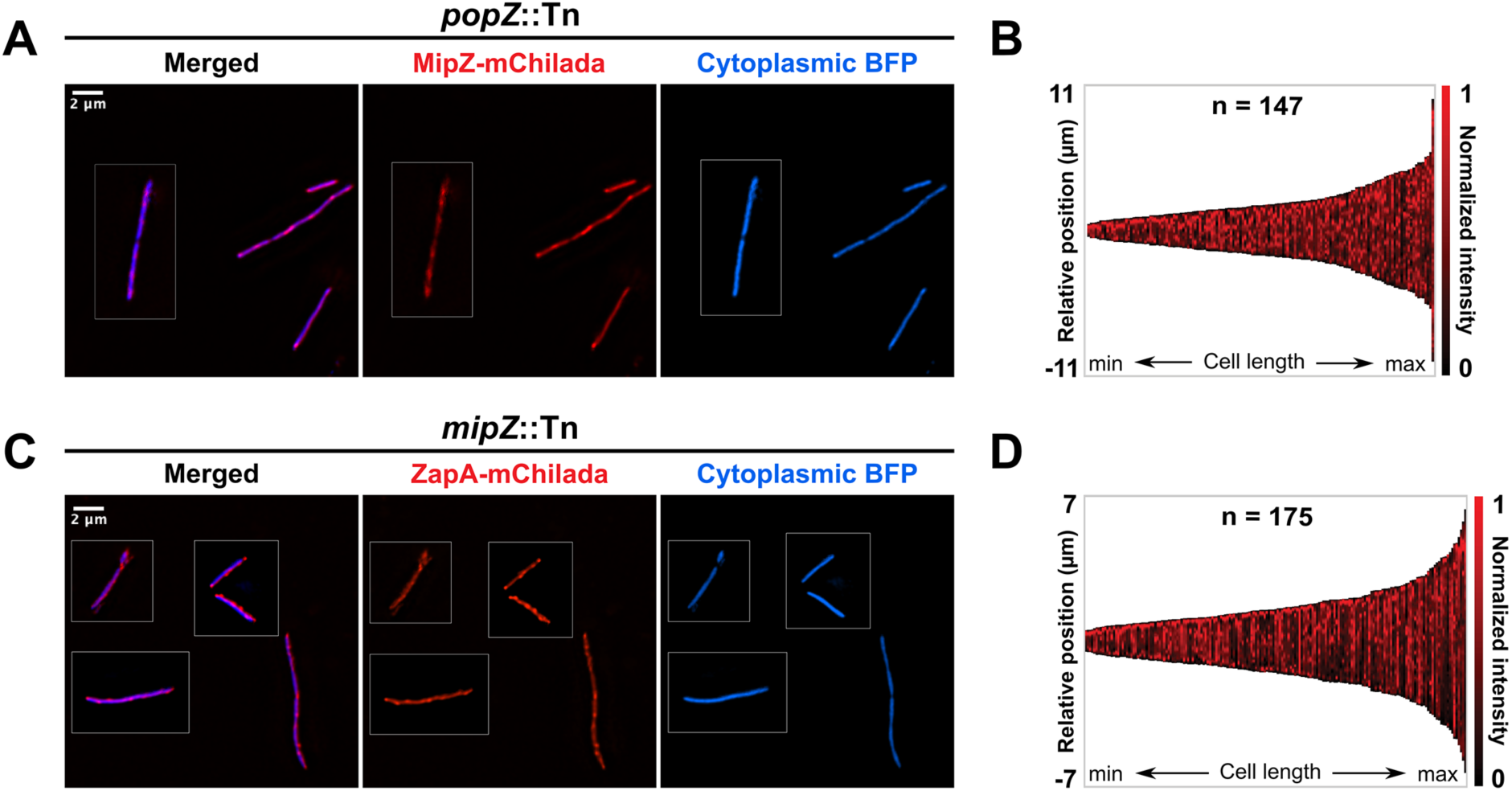
MipZ and ZapA are mislocalized in *popZ*::Tn and *mipZ*::Tn mutants, respectively. Representative epifluorescence micrographs and demographs of live *R. parkeri* grown in A549 host cells at 24 hpi. **A-B**. *R. parkeri popZ*::Tn expressing *mipZ-mChilada* from the *zapA* promoter (EGRp0098). **C-D.** *R. parkeri mipZ*:Tn expressing *zapA-mChilada* from the *zapA* promoter (EGRp0129). *R. parkeri* also constitutively expressed cytoplasmic *tagBFP* from the *ompA* promoter. TagBFP signal was used to delineate cell outlines for plotting mChilada signal across cell length to construct the demographs in MicrobeJ, which represent a cell population from one of three technical replicates for each strain. Boxed images represent different fields of view from combined to display more representative cells.

### Consequence of dysregulation in chromosome organization and cell division on *R. parkeri* fitness

To characterize the functional ramifications of perturbations in chromosome segregation and cell division in the context of intracellular growth and cell-to-cell spread, we performed plaque size measurements for each *R. parkeri* mutant defective in one component of these pathways. Plaques are zones of dead and lysed cells caused by *R. parkeri* growth and/or spread in the host cell monolayer. The size of the plaque reflects efficiency of growth, intracellular survival, and cell-to-cell spread of a given strain. While the plaque size of the *mipZ*::Tn mutant strain was comparable to WT, loss of function of *parA or parB* elicited a significant reduction plaque size (**Figure 6**). This small plaque phenotype aligns with the original *R. parkeri* screen from which the *parB*::Tn strain was isolated based on its reduced plaque size [7]. Surprisingly, *popZ*::Tn showed a slightly larger plaque size than WT *R. parkeri* (**Figure 6**), implying the possibility of an unknown fitness advantage acquired through loss of polar chromosome anchoring or other activities mediated by PopZ. We note that a *rarp-1::*Tn mutant was reported to yield a small plaque [36], suggesting that our *popZ* transposon insertion does not disrupt expression of the downstream *rarp-1* gene. Because all the mutants tested have similar cell division defects and morphology, these data implicate chromosome segregation by ParABS, specifically, as critical for growth and/or spread in a monolayer of host cells.

**Figure 6.**
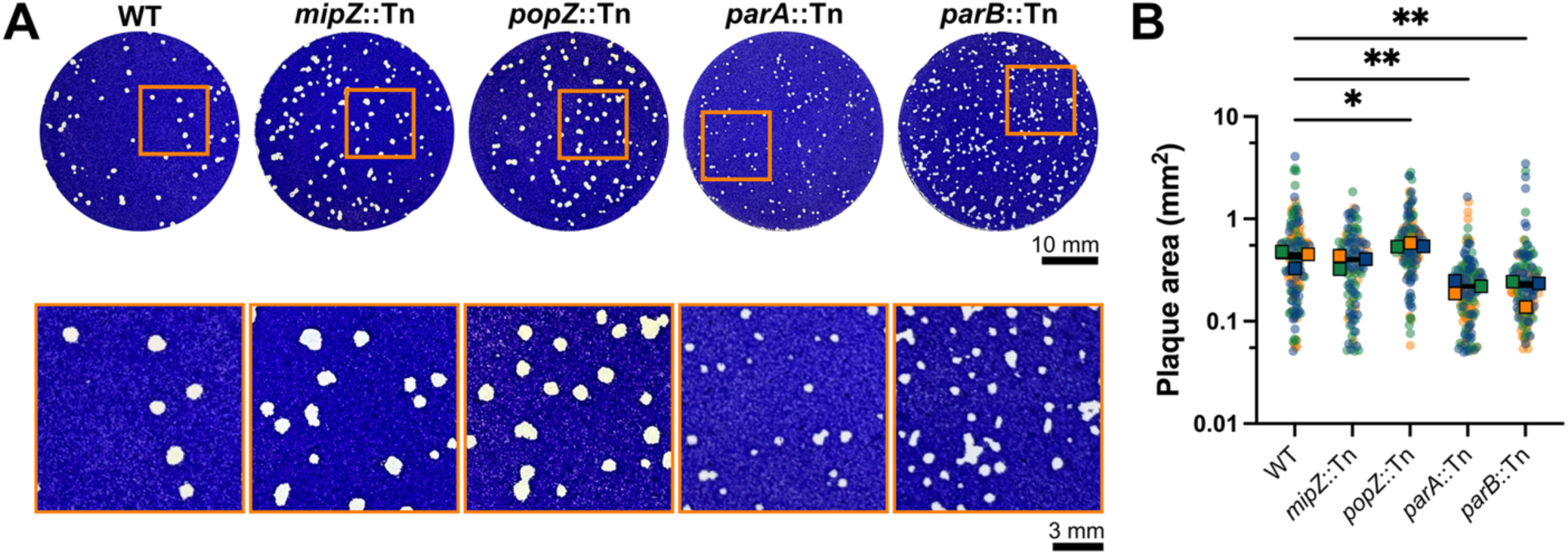
Perturbations to genome organization impact *R. parkeri* fitness in a monolayer of mammalian host cells. **A**. Representative images of plaques in Vero cell monolayers infected with indicated *R. parkeri* transposon mutant strains and counterstained with crystal violet at 7 days post-infection (dpi). Areas bounded by orange boxes were enlarged. *mipZ*::Tn = EGRp0022; *popZ*::Tn = EGRp0128; *parA*::Tn = EGRp0052; *parB*::Tn = EGRp0034. **B.** Quantification of plaque areas from (A). Circles represent individual plaque sizes from three independent technical replicates (blue, orange, and green [N=3]; 50 plaques/replicate [n=50]) and outlined boxes represent median values for each replicate. Lines indicate the mean of replicate medians. A one-way ANOVA with Dunnett’s multiple comparisons test was performed to compare population median values (N=3) for each transposon mutant to WT *R. parkeri* and to determine *P* values (*, *p* = 0.05; **, *p* < 0.01), assuming a Gaussian distribution of median plaque size.

## DISCUSSION

Rickettsiae are biomedically important human pathogens, but their intimate connection with the host intracellular niche has substantially slowed progress in understanding their cell biology [9]. In this study, we applied recent innovations in imaging [9] and genetics [7,8] to investigate the cellular organization of the SFG model *R. parkeri* with specific focus on chromosome segregation dynamics and division plane placement. We demonstrate that the chromosome is likely organized in a polar fashion that is leveraged to position the division site at the right time and place. Chromosome segregation, specifically, is important for *R. parkeri’*s intracellular growth and spread within a monolayer of host cells. This work illustrates for the first time functionally significant intracellular polarity in a *Rickettsia* species, with consequences for its fundamental biology and, perhaps, host interactions.

Our data showed that chromosome dynamics and spatial control of cytokinesis are mediated by the PopZ-ParABS-MipZ axis in *R. parkeri*, consistent with the paradigm established in *C. crescentus*. On the other hand, the master regulator of cell polarity PopZ, which forms a polar condensate, occupies a single cell pole for the entire cell cycle in *R. parkeri* (**Figure 3E-F)**, but still physically engages with ParA and ParB (**Figure 4**). PopZ is exclusively encoded in Alphaproteobacteria and displays diverse subcellular localization patterns in addition to the unipolar-to-bipolar dynamics in *C. crescentus*. In the plant pathogen *Agrobacterium tumefaciens* [37] and the mammalian pathogen *Brucella abortus* [38], known for their unipolar growth, PopZ resides at the actively growing pole and moves to midcell during division. After cell division, it marks the new growth pole of each daughter derived from the division site. In the stalked, budding bacterium *Hyphomonas neptunium*, PopZ initially localizes to the old pole before it relocates to the opposite pole in the nascent bud to mediate ParB-*parS* polar docking after chromosome translocation [39].

We cannot yet monitor the cell cycle by timelapse microscopy in *R. parkeri*, so the identity of the pole(s) marked by PopZ, or whether PopZ changes its polar location during cell cycle progression, remains unknown. We suggest two possible models for PopZ localization in *R. parkeri*. In the static model, PopZ remains at a single pole in the mother cell and forms a polar condensate via *de novo* biosynthesis in the progeny. In the dynamic model, PopZ relocates from the old pole to the new pole during chromosome segregation like in *H. neptunium* [39]. Another possibility is that the two models are connected: PopZ dynamically moves from the old pole to the new pole in the mother cell, while accumulating anew either at the nascent pole or the old pole in the newborn offspring.

Prior discoveries on the regulation of PopZ condensate dynamics lead us to favor the static model (**Figure 7**). First, in *C. crescentus*, the polar organelle development protein PodJ plays a primary role in triggering the dynamic accumulation of PopZ at the new pole, as PopZ in *C. crescentus* cells without PodJ failed to adopt a bipolar pattern at the predivisional stage [40,41]. PopZ-PodJ direct interaction is conserved in *A. tumefaciens* and *H. neptunium*, both of which exhibit cell cycle-dependent PopZ dynamics [42]. Rickettsiales, however, lack a PodJ homolog (**Figure 1**), suggesting PopZ in these species might not undergo dynamic re-localization during the cell cycle. Moreover, PopZ comprises three domains: an N-terminal client binding region, an intrinsically disordered region (IDR) tuning its material properties, and C-terminal helices mediating PopZ oligomerization and condensation [19,20,34–36,38,46,47]. Notably, despite conservation of the overall protein architecture, the IDR in *R. parkeri* PopZ is severely truncated compared to most Alphaproteobacteria, suggesting distinct condensate dynamics and biophysical properties. Specifically, shortening the IDR renders *C. crescentus* PopZ more dense and less mobile [33], so a truncated IDR in *R. parkeri* PopZ might similarly reduce condensate dynamics. We hypothesize that reduced condensate dynamics and absence of PodJ support stable accumulation of PopZ at one *R. parkeri* cell pole, with a new condensate formed *de novo* in the daughter that fails to inherit one. Whether there is a preference for assembly of a new condensate at the new or old pole is not clear.

**Figure 7.**
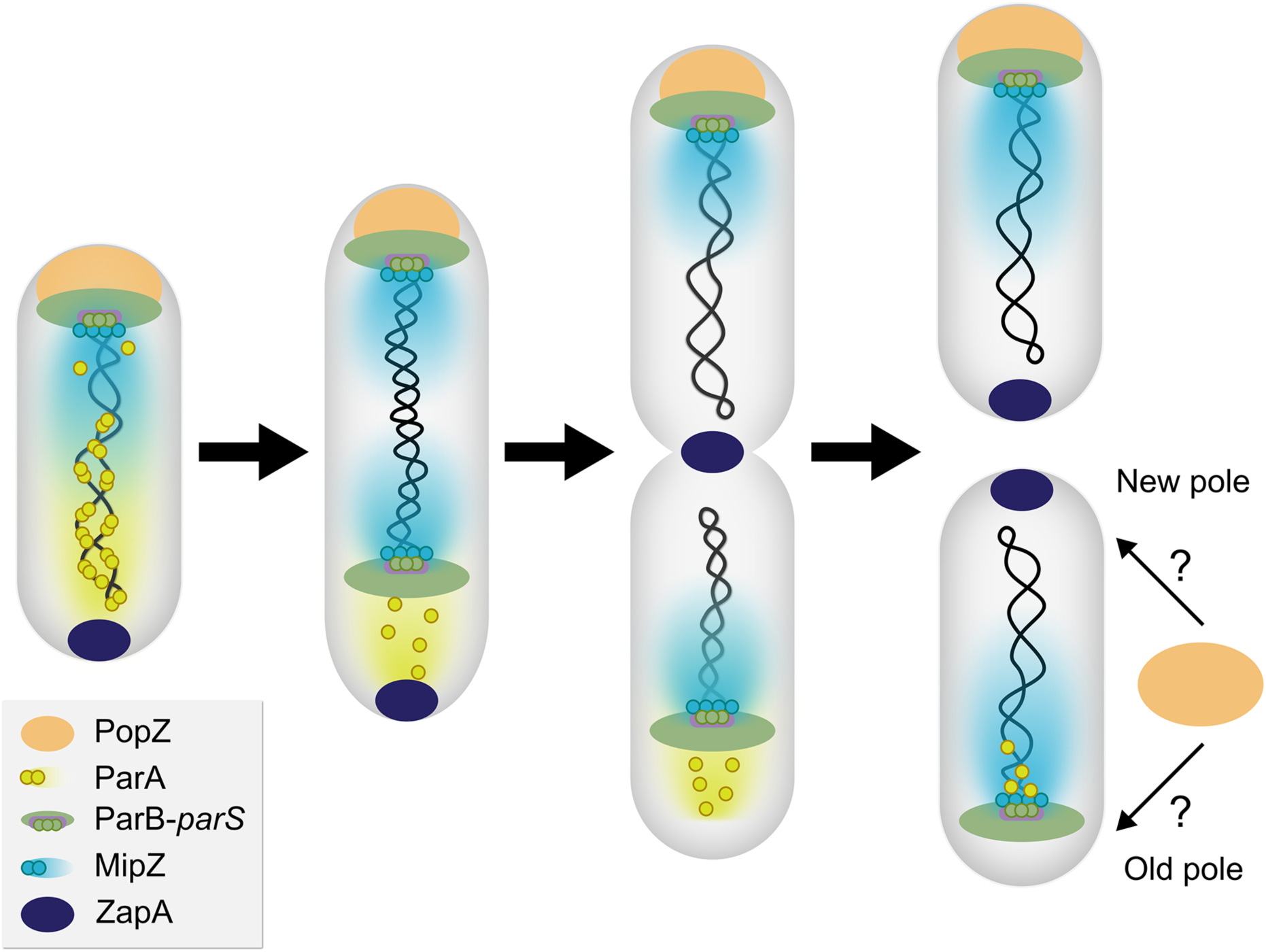
Model for the coordination of chromosome segregation and cell division in *R. parkeri.* PopZ (orange) self-assembles into a condensate and directly binds to ParB-*parS* (green and purple) to tether the centromere at one cell pole in G1. ParA dimers (yellow) coat the genome and dissociate from the DNA as monomers upon contact with ParB. Upon entry to S phase, *R. parkeri* duplicates the chromosome, and a second ParB-*parS* complex moves across the cell in the wake of the receding ParA gradient. The migration of the second ParB-*parS* copy allows ParB’s direct interactor, the division antagonist MipZ (light blue), to establish a bipolar gradient that is highest at the cell poles. As a result, the middle of the cell with the lowest MipZ concentration is the only permissive zone for cell division (marked by ZapA, dark blue) to occur. The subcellular localization dynamics of these factors aligns with the model of chromosome segregation and division plane placement in *C. crescentus*, except that *R. parkeri* PopZ remains unipolar across the cell cycle, raising questions about PopZ condensate dynamics, pole selectivity, and polar chromosome anchoring in the non-PopZ pole in *R. parkeri*.

Regardless of which PopZ model holds true, ParB-mNG exhibited a clear bipolar pattern. In the model for chromosome segregation in *C. crescentus*, the PopZ focus formed at the new pole captures the migrating ParB-*parS* complex to secure the second centromere copy to that pole [18,19]. In *R. parkeri*, we speculate three fates for the attachment of the segregated ParB-*parS*: (1) it could be tethered at the new pole by PopZ that either forms *de novo* (as in the static model proposed in **Figure 7**) or switches poles during chromosome replication (as in the dynamic model), (2) its attachment at the PopZ-free pole might occur in a PopZ-independent manner, or (3) attachment at the new pole is not required. The fact that *R. parkeri* PopZ retains direct interaction with ParA and ParB (**Figure 4**) and is required for proper regulation of cell division via MipZ (**Figure 5A**) underscores the intimate interplay between chromosome partitioning, centromere anchoring, and spatial positioning of the division septum across distant bacterial species.

In *C. crescentus*, *parA* and *parB* are essential for viability [45,46]. In contrast, the isolation of *parA* and *parB* transposon mutants indicates that neither protein is absolutely required for *R. parkeri* survival. The dispensability of ParA and ParB implies the existence of other factors that partially compensate for their loss in organizing the genome. The structural maintenance of chromosomes (SMC) protein complex is a widely conserved regulator for bacterial nucleoid compaction, organization, and segregation [47–56]. Crucially, SMC plays a major role in chromosome segregation in bacterial species missing ParA, such as *Streptococcus pneumoniae* [57] and *Staphylococcus aureus* [58], or completely lacking the ParAB system, like *E. coli* [59–62]. Strikingly, *R. parkeri* and other Rickettsiales lack SMC homologs, implicating other contributing factors. Likely candidates include nucleoid-associated proteins that are encoded in *R. parkeri* including HU family proteins [63], EbfC/YbaB [64], and/or GapR [65]. The lack of SMC and the non-essentiality of ParAB is surprising and raises the question of how chromosomal architecture, organization, and dynamics are orchestrated in *R. parkeri*.

Unlike *popZ*::Tn and *mipZ*::Tn, the *parA*::Tn and *parB*::Tn mutants exhibited a markedly smaller plaque size compared to WT *R. parkeri* (**Figure 6**). Alterations in plaque size could reflect defects in intracellular proliferation, immune evasion, motility, and/or cell-to-cell spread. Although each of these mutants leads to cell division defects, *parA*::Tn and *parB*::Tn are unique in that they likely fail to segregate the chromosome, leading to global disruption of the spatial organization of the genome. The nucleoid has been proposed as a key spatial regulator of bacterial cells, which typically lack membrane-bound organelles or long-range cytoskeletal systems [66]. Indeed, the nucleoid serves as a platform for self-organizing segregation systems [67], and gene location can dictate localization of the encoded protein [68,69]. We hypothesize that the disruptions to genome organization that result from loss of ParA or ParB have broad effects on cellular organization impacting everything from fundamental processes to host interactions.

Overall, our data provide insights into the cell cycle dynamics of chromosome segregation in coordination with spatial selection of the division plane in *R. parkeri* that are likely to apply to other rickettsial pathogens. These previously understudied yet crucial aspects of rickettsial biology define unappreciated intracellular polarity that plays a critical role in fundamental cellular processes. Cell polarity is further implicated in replication, survival, and/or spread during infection of host cells and is, thus, an important topic for future studies. This work highlights the utility of our quantitative imaging approaches and transposon mutant library in uncovering critical cell biological processes in obligate intracellular pathogens.

## MATERIALS AND METHODS

### Cell culture

A549 human lung epithelial and Vero monkey kidney epithelial cell lines were obtained from the American Type Culture Collection. Cells were cultured at 37 °C, 5% CO_2_ with 90% relative humidity. A549 cells were grown in DMEM (Life Technologies) supplemented with high glucose (4.5 g/l), L-glutamine, and 10% heat-inactivated fetal bovine serum (Hi-FBS). FBS was heat-inactivated by placing the FBS (Atlas Biologicals) in a water bath at ∼56 °C for 30 min. Vero cells were grown in DMEM (Life Technologies) supplemented with high glucose (4.5 g/l), L-glutamine, and 5% Hi-FBS. All eukaryotic cell lines were thawed and passaged ∼10 times until stable, growth rates were established before use and discarded after ∼50 passages.

### Plasmids and strains construction

All plasmids, oligos, and cloning strategies are listed in **Supplemental Table S1**. Plasmids transformed into *R. parkeri* are derived from pRAM18dA [70,71] with genes encoding antibiotic resistance (*rifR* and *specR*) and cytoplasmic fluorescent proteins (*tagBFP*, *gfpuv*, *aausFP1*) placed under the *ompA* promoter with the *ompA* terminator sequence. Strains used are listed in **Supplemental Table S2**.

### Transposon mutagenesis

The *parB::Tn* mutant was a gift of Matthew Welch (UC-Berkeley) [7]. Himar-1-based transposon mutagenesis of *R. parkeri* to generate the *mipZ::Tn, popZ::*Tn, and *parA::Tn* mutants was conducted using pMW1650, pEGTn01, or pEGTn02, respectively, as previously described [72]. pEGTn01 was generated as described for pEGTn02, but lacks a ∼500 bp insertion between the spectinomycin resistance and AausFP1 genes that was found to stabilize the plasmid in *Escherichia coli* (the stabilized plasmid is pEGTn02). The full sequence of pEGTn01 is in **Supplemental Figure S4**. To generate transposon mutants, freshly prepared or frozen electrocompetent *R. parkeri* were electroporated with ∼1 µg of the indicated transposon construct. After electroporation, the bacteria were added to a 6-well plate with confluent monolayers of Vero cells. Host cell infection was carried out at 37 °C for 30 min with rocking. After this period, an overlay of DMEM supplemented with 2% (v/v) heat-inactivated FBS and 0.5% (w/v) ultra-pure agarose was added to each well. The culture was then incubated at 33 °C, 5% CO_2_ overnight. The following day, a 1 mL overlay of DMEM + agarose containing 200 ng/ml rifampicin (pMW1650) or 50 µg/mL spectinomycin (pEGTn01 or pEGTn02) was added. The culture was allowed to incubate at 33 °C, 5% CO_2_ until plaques formed.

Plaques were picked, resuspended in 200 µL of BHI broth, and frozen at −80 °C until processing. For expansion, 100 µL of the resuspended plaque was used to infect one well of a 6-well plate containing confluent monolayers of Vero cells; spectinomycin at 50 µg/mL was maintained in the medium. Once the infection reached completion, the bacteria were harvested. Bacteria were stored at −80 °C in BHI, but a small sample was retained for PCR mapping of the transposon insertion site.

Semi-random PCR was conducted on boiled bacterial lysate from the expanded *R. parkeri* transposon mutant. The first round of PCR used specific primers for the transposon sequence (oEG1853 (*mipZ*::Tn) or oEG1941 (*popZ*::Tn and *parA*::Tn)) and a universal primer (oEG1854 (*mipZ*::Tn and *popZ*::Tn) or oEG2015 (*parA*::Tn)). The universal primer contained a specific G/C-rich 20-nt, 5’ anchor, a random 10-nt sequence, and a 3’ pentamer. The products from the first round of PCR were purified using the Monarch Spin PCR and DNA Cleanup Kit for use in the next round of nested PCR. The second round of PCR used primers for the transposon (oEG1856 (*mipZ*::Tn) or oEG1942 (*popZ*::Tn and *parA*::Tn)) and the 5’ anchor of the universal primer (oEG1857 (*mipZ*::Tn and *popZ*::Tn) or oEG2016 (*parA*::Tn)). The amplified DNA was purified and sent for Sanger sequencing. Transposon mutants were each subjected to whole-genome sequencing to confirm transposon insertion sites and ensure no other mutations were present in the genome.

### *R. parkeri* transformation

*R. parkeri* were transformed using a small-scale electroporation protocol as described previously [9]. Vero cells were grown to confluency in a T175 cm^2^ flask. Media were aspirated, and cells were washed with ∼10 mL of phosphate-buffered saline (PBS). 100 µL of WT or transposon mutant *R. parkeri* diluted in 3 mL of brain heart infusion (BHI) broth was added to the Vero cells. The infected flask was placed in a secondary container and incubated for 30 min at 37 °C in a waving shaker. Upon infection, eukaryotic cells were incubated in DMEM (high glucose, +L-glutamine) with 2% Hi-FBS and either 200 ng/ml rifampicin or 50 µg/mL spectinomycin for the corresponding transposon mutant. No antibiotic was added to the media of Vero cells infected with WT *R. parkeri*.

After ∼4 days, when ∼80% of Vero cells started to round and lyse as a result of infection, cells were scraped into the growth media, collected in 50 mL conical tubes, and centrifuged at 1,700 x g for 5 min at 4 °C in a JS-75 rotor. Infected Vero cell pellet was resuspended in 3 mL of cold K-36 buffer (0.05 M KH_2_PO_4_, 0.05 M K_2_HPO_4_, pH 7, 100 mM KCl, and 15 mM NaCl). Intracellular *R. parkeri* was released by mechanical bead disruption previously described [7], where the homogenate was transferred into a 15 mL conical tube containing ∼2 g of 1 mm glass beads and vortexed on setting 7 using a Pulsing Vortex Mixer (Fisher Scientific Cat #02215375), with two alternating 30 s pulses immediately followed by 30 s ice incubations after each pulse to mechanically lyse the infected eukaryotic cells. Host cell debris was removed by filtering the homogenate through a 2 µm filter (Neta Scientific). The filtered homogenate was transferred to two 1.5 mL tubes and centrifuged at max speed (16,300 x g) for 2 min to pellet *R. parkeri*.

To prepare electrocompetent *R. parkeri*, the bacterial pellet was washed three times by resuspending in 1 mL of cold, filter-sterilized 250 mM sucrose and centrifuged at max speed (16,300 xg) for 2 min. After the third wash, the bacterial pellet was resuspended in 50 mL of cold, filter-sterilized 250 mM sucrose mixed with 6-10 µg of DNA, and electroporated at 1.8 kV, 200 Ω, 25 μF, 5 ms in a prechilled, 0.1-cm-gap cuvette using a MicroPulser (Bio-Rad). Electroporated bacteria were immediately recovered in 600 µl of cold BHI. Next, 100 µL of electroporated bacteria was diluted in 200 µL of BHI and added to each well of confluent Vero in a six-well plate after the media was aspirated and cells were washed with ∼1 mL of PBS. After 30-min incubation at 37 °C in a waving shaker, infected Vero cells were overlaid with DMEM (high glucose, +L-glutamine) supplemented with 2% Hi-FBS and 0.5% (w/v) ultra-pure agarose (Invitrogen; 16-500) and were incubated at 33 °C, 5% CO_2_ in a high-humidity chamber. To select for positive transformants, a second agarose overlay containing DMEM (high glucose, +L-glutamine) with 2% Hi-FBS, 0.5% (w/v) ultra-pure agarose (Invitrogen; 16-500) and 200 ng/mL of rifampicin, or 50 µg/mL of spectinomycin, or both, was added 16 – 24 hpi. The plates were incubated at 33°C, 5% CO_2_ for 5–10 days in a high-humidity chamber before plaque selection (see below).

### *R. parkeri* standard strain expansion, preparation, and enumeration

After ∼5-10 days after electroporation, ∼6-10 visible plaques were collected, resuspended in 200 µL of BHI and stored at −80 °C. To expand confirmed clones producing fluorescent signal, the 200 µL of plaque resuspension was split into two infections: one 100 µL of plaque suspension was used to infect one well of confluent Vero monolayer in a six-well plate and the other 100 µL was used to simultaneously infect a well of confluent Vero in a second 6-well plate (MatTek; no. 1.5 coverslip, 20 mm glass diameter, uncoated plates, part no. P06G-1.5-20-F) for imaging-based screening. Infected Vero cells were incubated in DMEM and 2% FBS supplemented with the appropriate antibiotics for ∼4-10 days, or when ∼80% of host cells rounding/lysis was observed. Transformed *R. parkeri* was isolated by the previously described mechanical bead disruption method, and bacterial pellet harvested from each well was resuspended in 200 µL of BHI broth and stored at −80 °C (passage 1). For further strain expansion, 100 µL of bacterial suspension was added to 3 mL of BHI broth and inoculated into one T175 cm^2^ flask of confluent Vero cells as described above. After ∼4-10 days, bacteria were harvested, resuspended in 1mL of BHI broth, aliquoted into 100 µL, and stored at −80 °C (passage 2). All experiments were conducted on bacterial aliquots harvested from passage 2 and freeze-thaw cycles were limited to one time per stock.

To enumerate bacteria, one aliquot of the expanded strain was serially diluted in BHI and inoculated into confluent Vero cells in a 6-well plate. After 30 min incubation at 37 °C in a waving shaker, an overlay of DMEM (high glucose, +L-glutamine) with 2% Hi-FBS and 0.5% (w/v) ultra-pure agarose (Invitrogen; 16-500) with appropriate antibiotics was added to each well. Plates containing infected Vero cells overlaid with DMEM agarose were stored at 33 °C, 5% CO_2_ in a high-humidity chamber for 7 days before plaques were visualized and enumerated to determine the PFU/ml.

### Epifluorescence microscopy and quantitative image analyses

Epifluorescence microscopy was performed on live, intracellular bacteria expressing cytoplasmic GFPuv, AausFP1, BFP and/or fluorescently labeled proteins in A549 cells. A549 cells were seeded in 24-well MatTek dishes (no. 1.5 uncoated coverslip, 13 mm glass, cat no.) and grown for ∼ 24 hrs to allow for formation of a confluent monolayer. Confluent A549 cells were infected with ∼ 33.3 µL of *R. parkeri* strains in a total volume of 100 uL of BHI by shaking at 37 °C for 30 min as described above. Infected cells were incubated in DMEM (high glucose, +L-glutamine) supplemented with 10% Hi-FBS without antibiotics at 33°C, 5% CO_2_ in a high-humidity chamber for ∼ 24 hours to allow intracellular bacteria to enter exponential phase. After ∼24 hours post-infection, media of infected A549 cells were aspirated, and cells were washed with ∼1 mL PBS, and infections were incubated in Fluorobrite^TM^ DMEM (high glucose, +L-glutamine) (Life Technologies) supplemented with 10% Hi-FBS (Atlas Biologicals) to enhance live-cell fluorescent signals. Z-stack image acquisition was conducted by live epifluorescence microscopy as previously described in [9]. Briefly, MatTek dishes were placed on an on-stage incubation chamber (Tokai Hit) or in an STX Stage Top Incubator (Tokai Hit), with the top lid temperature set at 56 °C, the chamber temperature at 33 °C, and CO_2_ at 5% for the duration of the experiment to maintain the host cells and bacterial viability. A Nikon Eclipse Ti inverted microscope (RRID:SCR_021242) was used to image samples through a Nikon Plan Fluor 100× (numeric aperture, 1.30) oil Ph3 objective, with a Photometrics CoolSNAP HQ2 cooled charge-coupled-device (CCD) camera and Nikon Elements Imaging Software. For fluorescence, GFP and AausFP1 signals were detected through an ET-GFP filter cube, BFP through ET-BFP, and mChilada through ET-dsRed.

Quantitative assessment of *R. parkeri* cell morphology and subcellular localization of fluorescent fusion proteins was conducted on images of three technical replicates of live *R. parkeri* using the FIJI [73] plug-in MicrobeJ [74] to generate fluorescence demographs and SuperPlots of cell length. To exclude out-of-focus bacterial cells, each Z-stack image was deconvolved before undergoing extended depth of field (EDF) processing using Nikon Elements Imaging Software, and bacterial cells undergoing motility during imaging were manually removed from the deconvolved-EDF processed images to ensure accurate analysis of individual bacterial cells in MicrobeJ. Cell outlines were identified by the cytoplasmic fluorescence signals alone or combined with the fluorescent fusion protein signals and were used to stratify bacterial cells by length. Demographs were created in MicrobeJ by plotting the profile of fluorescently tagged proteins signals as a function of cell length. To determine the median cell length, the cytoplasmic fluorescence cell profiles were quantitatively analyzed by MicrobeJ, and SuperPlots of cell length and their replicate mediates were plotted using GraphPad Prism [75].

To generate demographs of ParA-msfGFP, msfGFP-ParB colocalization with mCherry-PopZ in *E. coli*, single-stack images of bacterial cells were acquired. *E. coli* cell outlines were identified via phase-contrast images, and fluorescent signals of fusion proteins were plotted as a function of stratified cell length.

### *E. coli* co-expression assay

*E. coli* strain BL21(DE3) was transformed with plasmids for expression of mCherry-PopZ alone or together with ParA-msfGFP and msfGFP-ParB (Supplemental Table S1) and stored in 50% glycerol at –80 °C. Before the assay, each transformed strain was plated on Luria-Bertani (LB) agar supplemented with 0.1 mg/mL ampicillin, 0.03 mg/mL chloramphenicol, or both. A single colony of each strain was inoculated into LB broth containing 0.05 mg/mL of ampicillin and/or 0.02 mg/mL of chloramphenicol and grown overnight in a shaking incubator at 37 °C. Next, the cultures were diluted 1:100 and allowed to grow to an OD_600_ of ∼0.3 by shaking at 37 °C. Expression of fluorescent proteins from plasmids was induced by 0.1% L-arabinose for ParA-msfGFP and msfGFP-ParB, and 0.5 mM IPTG for mCherry-PopZ for an additional 2 hours of growth at 30 °C. Approximately 2 µL of *E. coli* cells was immobilized on 1% (w/v) ultra-pure agarose pads prior to imaging.

### Plaque size assay

Vero cells were grown in 6-well plates to confluency and washed with PBS. Purified *R. parkeri* was diluted in BHI broth to concentrations sufficient to yield approximately 20-200 plaques per well. Bacteria were added to the Vero cells and then incubated with rocking at 37 °C for 30 min. The infected monolayer was overlaid with DMEM supplemented with 2% FBS, 0.5% ultra-pure agarose (Invitrogen; 16-500), and appropriate antibiotics. The plates were incubated at 33 °C with 5% CO_2_ for 7 days. The agarose was then removed, and the cells were fixed with 95% ethanol and stained with crystal violet solution (2% crystal violet [w/v], 0.8% ammonium oxalate [w/v], and 20% ethanol [v/v]). Stained monolayers were washed with deionized water and dried. Each well was imaged, and plaque area was measured using ImageJ/Fiji. The known diameter of the wells was used to convert pixels to length.

### Statistical analysis, experimental variability, and reproducibility

All experiments were performed in technical triplicates. All statistical analyses were performed in GraphPad Prism 9 using ordinary one-way Analysis of Variance (ANOVA) with Tukey’s or Dunnett’s multiple comparisons test to compare parametric (Gaussian/normal) median cell length and plaque-forming unit areas distribution. *P* values below 0.05 were considered statistically significant.

## ACKNOWLEDGEMENTS

We would like to thank Rebecca Lamason (MIT), Matthew Welch (UC-Berkeley), Grant Bowman (U of Wyoming), and their laboratories for strains, reagents, and advice. We thank Dezmond Cole for sharing strains. Research reported in this publication was supported by the National Institute of Allergy and Infectious Diseases of the National Institutes of Health under Award Numbers R21AI96675 and R61AI79998 (to EDG). The content is solely the responsibility of the authors and does not necessarily represent the official views of the National Institutes of Health. This work was also supported by a Randall Reed Scholar Award from the Johns Hopkins University School of Medicine to EDG.

---

**Supplemental Figure S1.**
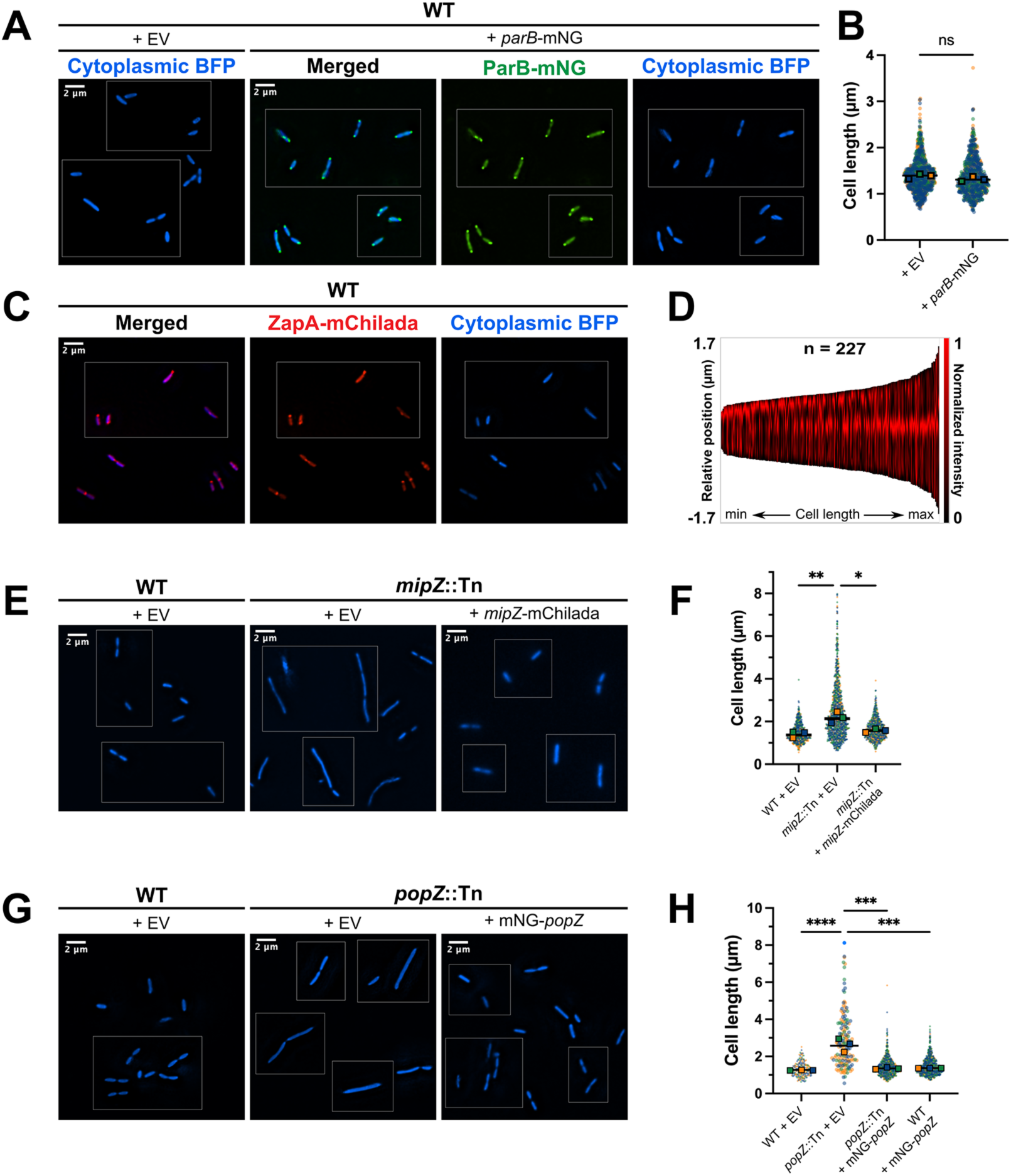
Fluorescent protein fusions preserve cell morphology and reveal the subcellular localization of cell division proteins in *R. parkeri*. **A-B.** Mild overexpression of ParB-mNG in WT *R. parkeri* does not affect cell morphology. A. Representative fluorescence micrographs of WT *R. parkeri* carrying an empty vector (EV) control (EGRp0039) and *parB-mNG* expressed under the *tolC* promoter (EGRp0015), both constitutively expressing cytoplasmic *tagBFP* driven by the *ompA* promoter in A549 host cells. B. Dot plot of cell length of cells captured in (A) using cytoplasmic TagBFP signal in MicrobeJ. Data are presented as in Figure 2B. Unpaired t-test assuming equal standard deviation for both strains and normal sample distribution was performed to compare the average population median values (n = 3) of the two strains and determine the *P* value (ns, not significant). **C-D.** ZapA-mChilada moves to midcell to mark division septum in constricting *R. parkeri* cells and a single new cell pole derived from the septum in newborn progeny. **C.** Representative fluorescence micrograph of live, intracellular WT *R. parkeri* expressing ZapA-mChilada from ZapA native promoter and constitutive cytoplasmic BFP from the OmpA promoter (EGRp003) in A549 host cells ∼ 24 hpi. **D.** Demograph plotted ZapA-mChilada signal as a function of cell length stratified by TagBFP signal in MicrobeJ using images of *R. parkeri* cells from (C). **E-H.**Fluorescence micrographs and cell length of live *R. parkeri* infecting A549 cells at 24 hpi. E-F. *mipZ*::Tn either expressing *mipZ-mChilada* from the *zapA* promoter and cytoplasmic *tagBFP* from the *ompA* promoter (EGRp0018), or EV control with just *tagBFP* (EGRp0081). Cytoplasmic TagBFP was used to identify cell outlines and compare cell lengths of *mipZ*::Tn strains with a WT *R. parkeri* carrying the EV (EGRp0002). G-H. *popZ*::Tn either expressing *mNG-popZ* from the *tolC* promoter and cytoplasmic *tagBFP* from the *ompA* promoter (EGRp0054), or carrying an EV with just *tagBFP* (EGRp0007). Cytoplasmic TagBFP signal was used to identify cell outlines and compare cell length of *popZ*:Tn strains with WT *R. parkeri* carrying the EV control (EGRp0039). Dot plots of cell length quantification in MicrobeJ using cytoplasmic TagBFP signal were presented similarly to Figure 2. Ordinary one-way ANOVA analysis with Tukey’s multiple comparison tests was performed to compare the population median values (n = 3) across *R. parkeri* strains and determine *P* values (*, p = 0.01, **, p < 0.01, ***, p = 0.0001, ****, p < 0.0001), assuming normal (Gaussian) distribution of the median length values.

**Supplemental Figure S2.**
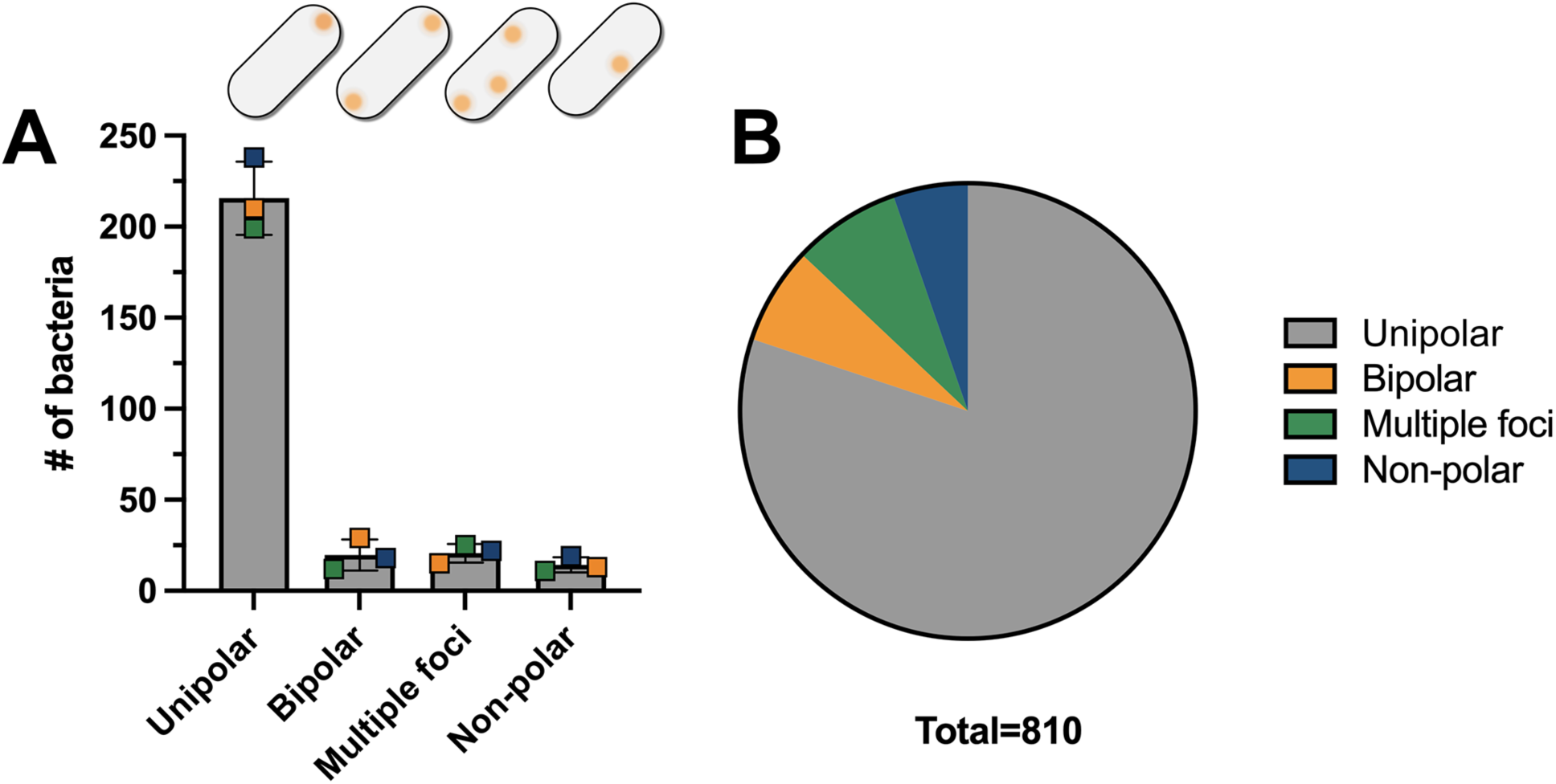
mNG-PopZ localizes mostly at one cell pole in *R. parkeri*. **A.** Bar graph showing the number of *R. parkeri* exhibiting the indicated subcellular localization patterns for mNG-PopZ captured in Figure 3E. Cells with no mNG-PopZ were not quantified. Outlined boxes represent the cell count for three technical replicates (blue, orange, and green) and error bars represent the standard deviation derived from the triplicates. **B.** Pie chart illustrating the percentage of the *R. parkeri* cells with different mNG-PopZ patterns from the combined number of cells from three technical replicates.

**Supplemental Figure S3.**
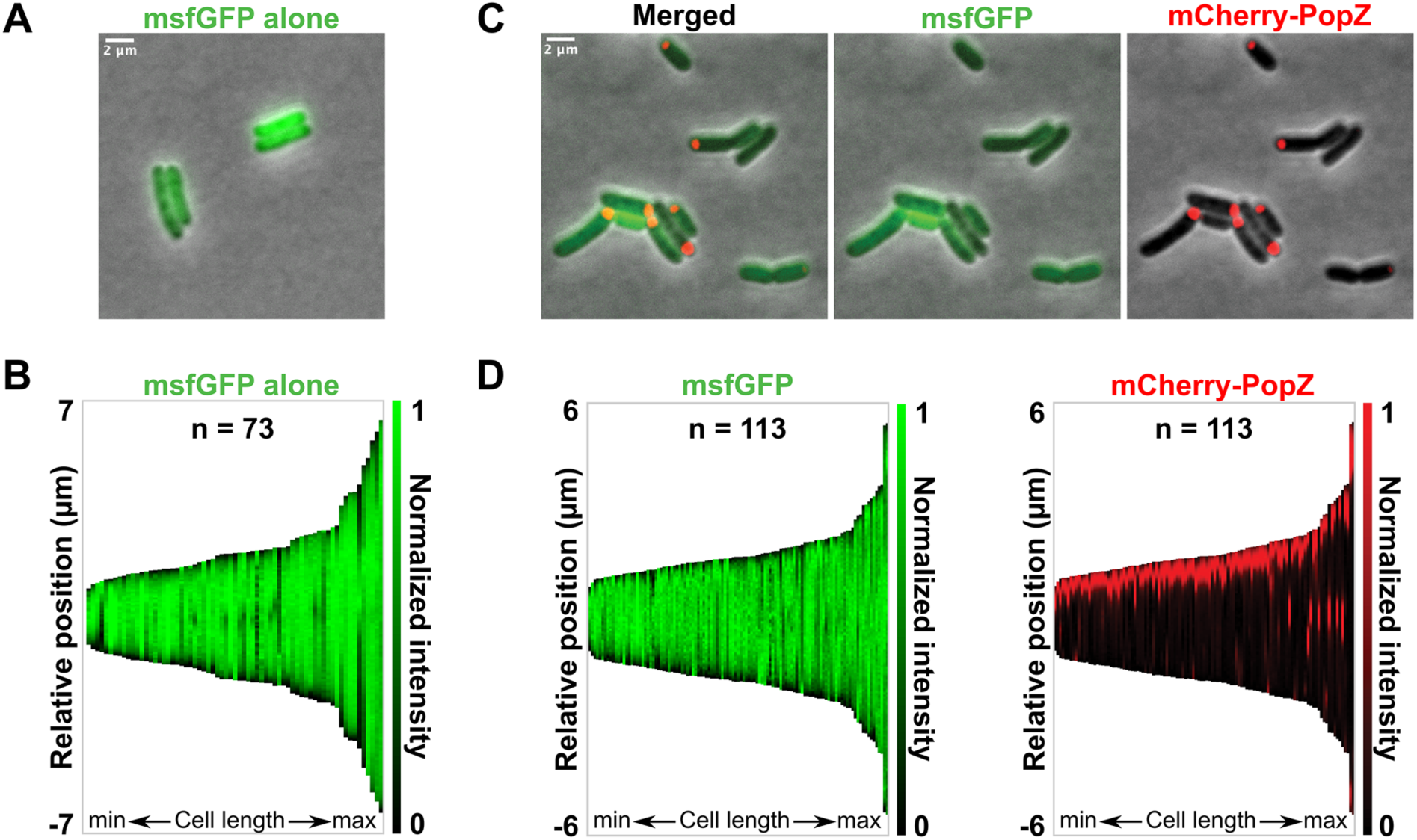
*R. parkeri* mCherry-PopZ forms a polar condensate and untagged msfGFP is diffuse in the cytoplasm when expressed in *E. coli*. Representative epifluorescence micrographs and demographs of *E. coli* BL21 cells expressing indicated fluorescent proteins captured after 2 hours post-induction. **A-B.** msGFP expressed alone was diffuse in *E. coli* (pEG2214). **C-D.** In the presence of *R. parkeri* mCherry-PopZ, which formed a condensate at one pole in *E. coli*, untagged msfGFP remained diffuse (pEG2215). Representative demographs of one of three technical replicates plotted msfGFP and mCherry signal as a function of cell length and oriented the cell pole with the strongest fluorescence to the top, similar to Figure 4.

**Supplemental Figure S4.**
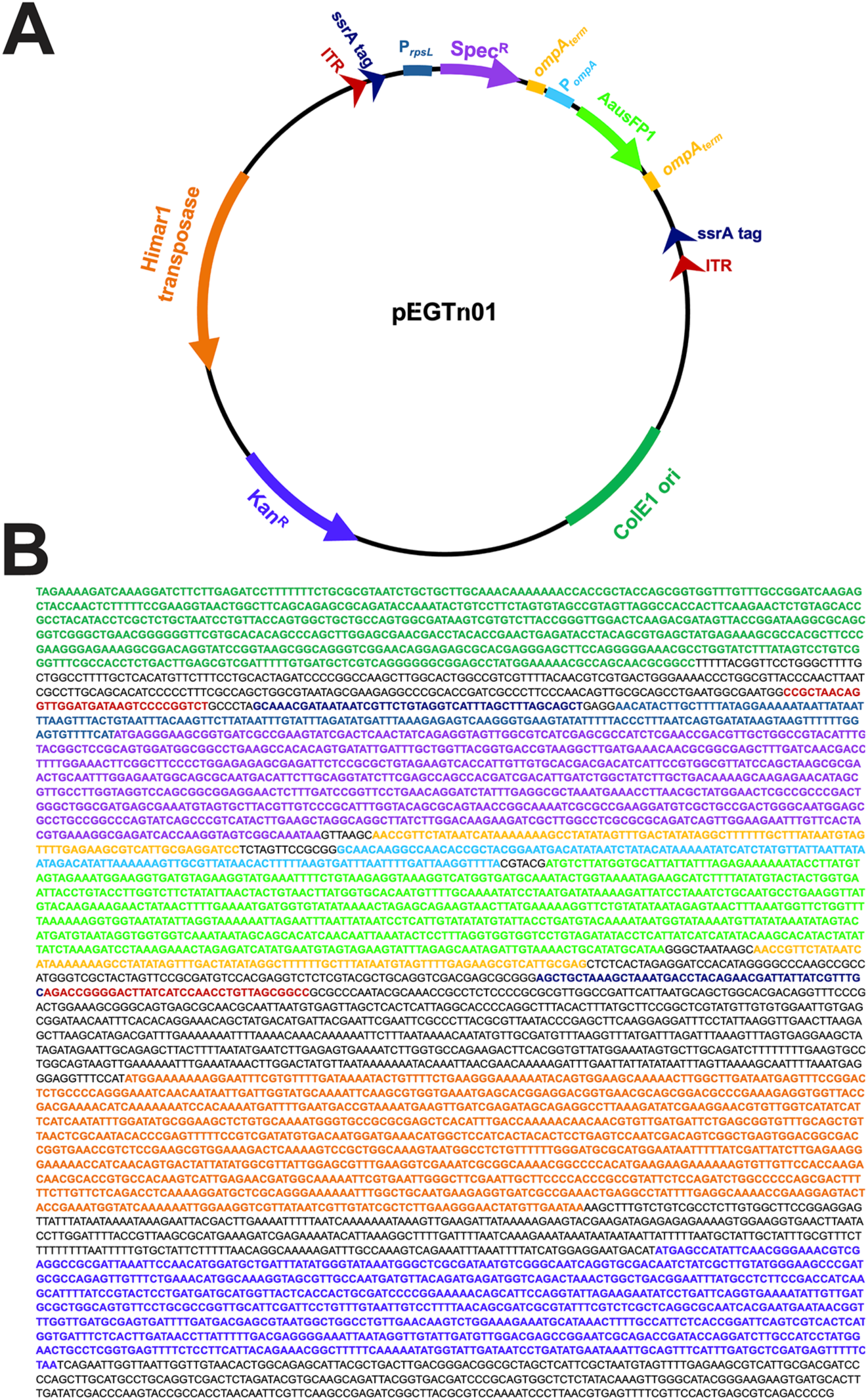
Plasmid map and sequence information of pEGTn01 transposon construct used to generate *R. parkeri* mutants. **A.** Plasmid map of pEGTn01. **B.** Color-coded featured sequences: ColE1 origin of replication and kanamycin resistance gene (to maintain in *E. coli*), spectinomycin resistance and AausFP1 gene (to select for *R. parkeri* transformants) under *rpsL* and *ompA* promoter, respectively, *ompA* terminator sequence, Himar1 transposase, inverted repeat sequence (ITR), and ssrA tag (to target protein product of in-framed insertion to proteolytic degradation).

Supplemental Table S1. Plasmids and oligos used in this study.

Supplementable Table S2. *R. parkeri* strains used in this study.

